# Statistical analysis provides new insights into patterns and mechanisms of the WANCY hotspot on vertebrate mitochondrial genomes

**DOI:** 10.64898/2026.10.16.759661

**Authors:** Chonghan Xu, Mark Wilkinson

**Affiliations:** Imperial College London, Department of Life Sciences, London-South Kensington, United Kingdom; Natural History Museum, Science Department, London-South Kensington, United Kingdom

**Keywords:** Mitochondrial genomes, Gene order rearrangement, Vertebrates, Convergent evolution, Mechanistic hotspot

## Abstract

The vertebrate mitochondrial genomes are commonly composed of 37 genes, and the order of these genes act as useful indicator of phylogenetic relationships. Tandem duplication and random loss (TDRL) is considered to be one of the most important mechanisms leading to mitochondrial gene order rearrangement in vertebrates. The WANCY region has been widely recognized as a TDRL hotspot. In the past, the lack of data limited the in-depth exploration of rearrangement patterns and mechanisms in hotspots. Through statistical analysis of a large amount of available NCBI sequencing data, combined with phylogenetic and bioinformatics methods, we obtained statistically significant data from the four classes of Amphibians, Reptiles, Mammalia and Osteichthyes. These data demonstrate three new convergent rearrangements in the WANCY region of vertebrates, which suggests that the frequency of convergent rearrangements in hotspots is higher than previously thought. We also compared the frequency of TDRL rearrangement in the WANCY region with two other similar small gene clusters (the IQM region and the HSL region). The results to a certain extent support the hypothesis that the WANCY region may be a mechanism hotspot. However, more molecular evidence and larger data volumes are needed to confirm the existence of mechanisms that cause rearrangement hotspots.

## 1. Introduction

Mitochondrial gene order is recognised as a useful indicator of phylogenetic relationships due to their conservativeness (Shi et al., 2015; Boore & Brown, 1998). Different gene orders are highly diagnostic in characterising groups of animals on different taxonomic levels (Boore & Brown, 1998; Boore, 2006). This point of view has been well applied in the Phylum Chordata, for example, diagnostic gene orders can be used for Marsupials and Crocodilians (Boore, 1999; Kumazawa & Nishida, 1995; Pääbo et al., 1991). The vertebrate mitochondrial genome is typically a 15-17 kb circular molecule containing the same 37 genes (Boore, 1999). These genomes exhibit different gene orders due to gene transposition (i.e., moving to a different position on the same strand), inversion (i.e., moving to the opposite strand but occupying the same position in the mitochondrial DNA), or both inversion and transposition (a combination of the first two events) (Sankoff et al., 1992; Kong et al., 2009; Gong et al., 2020). However, these mechanisms cannot explain the presence of pseudogenes and other intermediate state genes in the genome (San Mauro et al., 2006). Tandem duplication and random loss (TDRL) explains rearrangements through the duplication of a segment of the genome followed by the random loss of one of the duplicates (Moritz et al., 1987; Boore, 2000). According to this model, tandem duplication involves imprecise termination, strand slippage, and/or mismatches, which results in mitochondrial DNA replication errors (Pluciennik et al., 2000; Moritz & Brown, 1986; Moritz & Brown, 1987; Macey et al., 1997). As the evidence of an intermediate step of TDRL, pseudogenes and residues of tandemly repeated segments supports this model (San Mauro et al., 2006; Macey et al., 1998; Mueller & Boore, 2005). On the basis of TDRL, several derived mechanisms of gene order rearrangement have also been proposed. Dimer-mitogenome and non-random loss (DMNL) describes the non-random process of aggregation and loss of excess genes according to transcriptional polarity following tandem duplication of the entire mitochondrial genome (Lavrov et al., 2002). Double replications and random loss (DRRL) proposed that in some flounders, control regions (CRs) are replicated and translocated. Subsequently, double duplication of the mitochondrial genome is initiated sequentially from both CRs, resulting in the duplication of gene segments in between. Then, one of the redundant genes will be discarded randomly (Shi et al., 2014). Although various models have been proposed, the specific mechanisms by which gene order is altered are not fully understood (Babbucci et al., 2014). Among these models, TDRL is considered to be the major mechanism causing rearrangement events in vertebrate mitochondrial genomes and is widely accepted for explaining most of the existing gene order rearrangements (San Mauro et al., 2006; Pereira, 2000; Inoue et al., 2003; Arndt & Smith, 1998; Beckenbach, 2012).

The “WANCY” genomic region has widely been recognized as one of the most well-known hotspots of gene order rearrangements by TDRL (San Mauro et al., 2006; Montaña-Lozano et al., 2022). The “WANCY” is a cluster of five consecutive tRNA genes that locates between nicotinamide adenine dinucleotide dehydrogenase subunit 2 (ND2) and cytochrome coxidase subunit 1 (COX1) in most of the vertebrate mitochondrial genomes. The tRNA gene cluster includes ‘tRNA^Trp^’ (W), ‘tRNA^Ala^’ (A), ‘tRNA^Asn^’ (N), ‘tRNA^Cys^’ (C) and ‘tRNA^Tyr^’ (Y). The size of each small tRNA genes is around 70bp. In almost all the vertebrates, the origin of replication of light strand (O_L_) locates between tRNA^Asn^ and tRNA^Cys^ (San Mauro et al., 2006), which makes the WANCY a special genomic region both structurally and functionally. O_L_ is a stem-loop secondary structure which initiates the light strand replication (Seligmann, 2010). Some studies have shown that genes flanking this structure may be more likely to be duplicated (Boore & Brown, 1998; Dowton & Austin, 1999; Boore, 2000). Recognized mechanisms which lead to tandem duplication are thought to be slipped-strand mispairing during replication and illicit priming of replication by tRNA (Moritz & Brown, 1986; Cantatore et al., 1987; Seligmann et al., 2006). In mitochondrial genomes, these are thought to particularly involve stem-loop structures (San Mauro et al., 2006). A number of previous studies have suggested that O_L_ may be involved in the process of gene rearrangements (San Mauro et al., 2006; Macey et al., 1997), mutational gradient (Raina et al., 2005) and asymmetric nucleotide compositional deviations (Perna & Kocher, 1995).

The present of O_L_ seems to justify the mechanistic reason causing additional gene order rearrangements in the WANCY region. However, TDRL events are not reflective of rearrangement events. The greater number of genes involved in a duplication event, the greater the probability that the order of genes changes after random loss (formula 1). Therefore, the duplications happen in segments comprised by multiple small genes are more likely to be detected, which makes regions like the WANCY a potential epistemic hotspot of rearrangement (San Mauro et al., 2006).

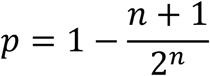

**Formula 1.** The probability of rearrangement after a single TDRL. N refers to the number of genes involved.

In addition to this, convergent rearrangement of gene order is also a subject of interest. Due to the rarity and complexity of gene order rearrangements, convergent evolutions may provide exceptionally useful data for phylogenetic inference (Macey et al. 1997; Boore & Brown 1998; Dowton et al., 2002). In the phylogenetic context, homogeneous gene orders can be used to characterise various evolutionary clades (Babbucci et al., 2014). In general, the chances of different taxa sharing the same rearranged gene order are thought to be very low (Babbucci et al., 2014). However, examples of convergent rearrangements occurring in different taxa do exist. For example, Siphonops species and Marsupials share the same WANCY region rearrangement (ACWNY) (Pabo et al. 1991; San Mauro, 2006). But so far, reports of convergence on hotspot regions of vertebrate mitochondrial genome rearrangements remain rare.

Over the past decades, insights into the patterns and mechanism of gene rearrangement in the mitochondrial genome have been limited by the amount of data available. In recent years, with the rapid development of sequencing techniques, the documented sequenced vertebrate genomes have exploded. As of December 2023, the number of complete vertebrate mitochondrial genomes recorded on GenBank has soared to 88128 (NCBI, 2023). Big data makes it possible to reveal rearrangement patterns and mechanisms from the statistical perspective. The aim of this study is to summarise existing sequenced data and investigate the WANCY genomic region, thus provide insights on the diversity of rearrangement patterns as well as the cause of the WANCY hotspot. To investigate the cause of the WANCY hotspot, we propose two competing hypotheses. The null hypothesis (H₀) is that WANCY does not have an intrinsically higher frequency of TDRL events than other comparable genomic regions. Instead, it appears as a hotspot because its cluster of five small genes increases the probability that a TDRL event will produce an observable gene order rearrangement (the epistemic explanation). Under this hypothesis, the observed rearrangement frequency in WANCY should be consistent with the expected frequency calculated from other tRNA gene clusters after accounting for differences in the probability of observable rearrangements. The alternative hypothesis (H₁) is that WANCY has an intrinsically higher frequency of TDRL events, potentially promoted by the presence of the light-strand origin of replication (OL) or other local molecular features (the mechanistic explanation). Under this hypothesis, WANCY should exhibit a higher frequency of independent TDRL rearrangements than expected from the background model, even after accounting for differences in rearrangement detectability.

In this study, we downloaded all the useable vertebrate mitochondrial genome records from NCBI GenBank, and then summarized the patterns of rearrangements. Multiple scripts and software were used to screen data and identify gene order rearrangement events caused by TDRL and thereby calculate the frequency of occurring TDRL in reality. This number was used to compare with the calculated frequency of TDRL in any other cluster of five small genes with no mechanisms promoting gene order rearrangement. This comparison has the potential to provide evidence to reveal if there is a mechanism which promotes additional gene order rearrangements in the WANCY region.

## 2. Methodology

### 2.1 Theory

We aim to compare the frequency of rearrangement of the WANCY region in reality (A%) with the frequency of rearrangement in any other cluster of five small genes (B%). However, for the consensus mitochondrial gene order of vertebrates, there’s no other cluster of five small gene.

Therefore, we focus on two different clusters of three consecutive tRNA genes each (IQM region and HSL region). The IQM region consists of ‘tRNA^Ile^’, ‘tRNA^Gln^’ and ‘tRNA^Met^’; The HSL region consists of ‘tRNA^His^’, ‘tRNA^Ser^’ and ‘tRNA^Leu^’. Then, B% can be obtained by using the frequency of occurring TDRL across the genome if there’s no mechanistic reason (C%) multiply the probability of occurring rearrangement after one time TDRL for five genes (D%). The rate of rearrangement in the IQM or the HSL region in reality (E%) divided by the possibility of rearrangement after one time TDRL for three genes (F%) is C%. D% and F% can be calculated via formula 1. A% and E% can be obtained via empirical data screening.

### 2.2 Data set construction

Firstly, the relatively complete mitochondrial genome files for a total number of 42391 (excluded humans) Mammalia, Reptiles, Amphibians, birds, Osteichthyes, Chondrichthyes were downloaded from NCBI GenBank (other vertebrate taxa were not analyzed due to the insufficient data quantity). All data were downloaded on 29^th^ Dec. 2023. Since the analysis needs only gene order annotations instead of nucleotide sequence, a Perl script from Xia, et al. (2016) was used to obtain gene names from the GenBank files. The Perl script obtains accession number, sequence length, species name, gene names in their original order, and total number of genes for each individual and generates a new txt file which will be used as the input file for the rearrangement identification script. A python script (SI_1.) was then used to resample the records. If there were multiple records for the same species, one was randomly retained and the rests were deleted, to ensure that each species occurs only once, thus avoiding sampling bias.

### 2.3 Rearrangement identification

Python scripts (SI_2.; SI_3.; SI_4.) were written to identify gene order rearrangements in the WANCY region, IQM region and HSL region for all individuals. We did not consider rearrangements that involve both the WANCY and other adjacent genes, since the calculation of the possibility of occurring rearrangements after one time TDRL needs fixed gene number.

We ignored a few cases which retain the duplicated sections in the genome, since the rearranged gene order can not be determined until the redundant parts are completely lost. Considering the following reasons, the positions of O_L_ (which recorded as “RO” on GenBank) were excluded. Firstly, the records on GenBank are not consistent, and the position of the O_L_ is not documented in a large portion of the data. Second, most of the current studies on convergent evolution only considered the positions of functional tRNAs. Finally, the TreeREx algorithm (will be introduced later) is unable to handle the case of inconsistent gene numbers. All identified rearrangements were exported to a txt file. Then, the rearranged genotypes of the rearranged species in each class were categorized to demonstrate the number of distinct gene orders generated.

### 2.4 Phylogenetic filtering

We performed phylogenetic filtering on all species in which gene order rearrangements have occurred to determine whether the rearranged gene orders were inherited from a common ancestor. These species that inherited genotypes from an ancestor were considered duplicates because they did not independently undergo a gene order rearrangement event.

Phylogenetic trees containing as many classified Mammalia (Upham et al., 2019), Reptiles (Pyron et al., 2013), Amphibians (Pyron & Wien, 2011; Portik et al., 2023), and Osteichthyes (Betancur-R et al., 2017) as possible were downloaded from the published papers. The ape package in R (Version 4.3.1) was used to prune these large phylogenetic trees to preserve the rearranged species as well as all related species along their evolutionary paths that has a determined gene order recorded on GenBank. These subtrees can be used to determine the inheritance relationships of these species.

TreeREx V1.85 (Bernt et al., 2008) was used to reconstruct the history of rearrangements. TreeREx is an algorithm based on an extended version of the CREx (Common Interval Rearrangement Explorer) algorithm (Bernt et al., 2007), which computes pairwise rearrangement scenarios for gene order data. It considers inversion, transposition, reverse transposition, and tandem repeat random loss (TDRL) scenarios (Bernt et al., 2008). TreeREx can automatically infer the mitochondrial gene order rearrangement events of real biological instances and thereby infer the evolutionary pathways leading to the observed gene order diversity (Bernt et al., 2008). We reconstructed the ancestral rearrangement patterns using the prunned subtree and a fasta file of the gene order of all species on the leaf tip as input files. Strong consistency method, weak consistency method and parsimonious weak consistency method were all considered to search for alternative rearrangements scenarios from a global perspective. When a rearrangement can be explained by both a single transposition or a single TDRL, we considered two separate scenarios, i.e., all such rearrangements were caused by transpositions; and all such rearrangements were caused by TDRLs. Finally, FigTree V1.4.4 was used to edit and visualize the phylogenetic tree and the rearrangement patterns.

### 2.5 Binomial test

Binomial test was conducted using R (Version 4.3.1) to determine whether the frequency of independent gene order rearrangement events which can be explained by TDRL in the WANCY region of vertebrates is significantly more than that of the IQM and the HSL region.

## 3. Results

### 3.1 Patterns of rearranged gene orders

Table 1 shows the number of individuals of different classes before and after resampling. The total number of vertebrates decreased from 42391 to 9419. 155 rearrangements involving only the WANCY region were determined. The rearranged gene orders are demonstrated in the following tables. According to table 2, table 3, table 4 and table 5, Osteichthyes showed the most rearranged gene orders in the WANCY region, in which 60 rearranged species displayed seven rearranged gene orders. The second most is Amphibians, among which displayed 6 rearranged gene orders. Reptiles and Mammalia both showed only one rearranged gene order. None rearranged gene order in the WANCY region, IQM region and HSL region was discovered in Birds and Chondrichthyes. In total, the 9419 vertebrates display 12 distinct gene orders in the WANCY region (one vertebrate consensus and 11 rearranged ones). According to table 6, among the six rearranged gene orders of Amphibians, three of them convergent to other classes or families. Seven rearranged gene orders of Osteichthyes displayed three convergent ones. The only rearranged gene order in both Reptiles and Mammalia convergent with species from other classes. Overall, four convergent gene orders were found in the recorded vertebrates. These convergences cover all the four classes where rearrangements were discovered. These identical or similar gene orders includes not only species from different classes, but also species from the same class but different orders or different families. For example, the four frog species *Trichobatrachus robustus*, *Hyperolius marmoratus*, *Hemisus marmoratu* and *Breviceps poweri* from different families showed the same gene order of ‘W’, ‘N’, ‘A’, ‘C’, ‘Y’ in the WANCY region. Although they all belong to the Anura order, they locate far apart on the phylogenetic tree. Moreover, the four Osteichthyes species *Sillago sinica*, *Sillago parvisquamis*, *Trachinops taeniatus* and *Benthalbella linguidens* which from three genus showed the same gene order in the WANCY region as in these frogs.

**Table 1.** Number of individuals by class.

|  | **TOTAL** | **TOTAL_Resampled** | **WANCY Rearrangements** |
| --- | --- | --- | --- |
| **Amphibians** | 1895 | 486 | 23 |
| **Mammalia** | 16991 | 1751 | 57 |
| **Osteichthyes** | 15044 | 4936 | 65 |
| **Reptiles** | 1830 | 661 | 10 |
| **Chondrichthyes** | 1672 | 257 | 0 |
| **Birds** | 4959 | 1328 | 0 |
| **Total** | 42391 | 9419 | 155 |

**Table 2.**
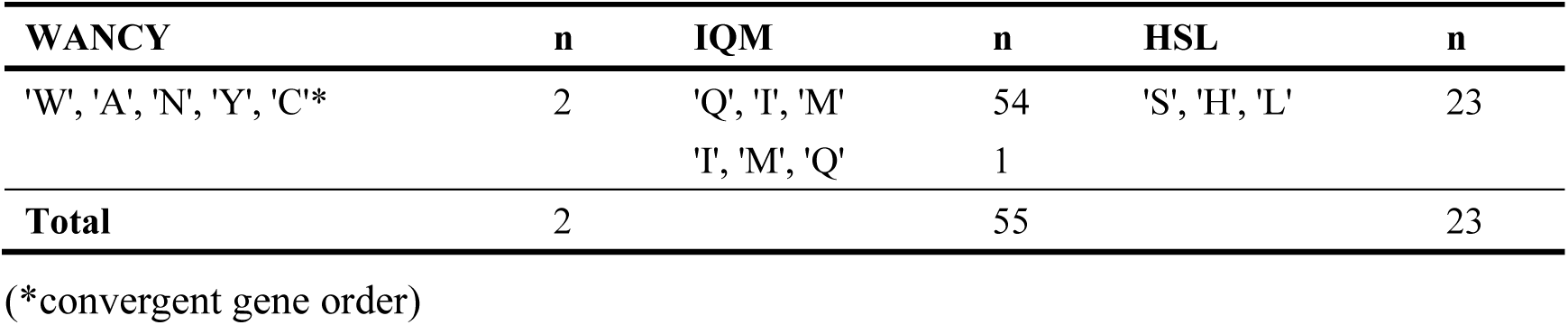
Patterns of rearranged gene orders of Reptiles.

**Table 3.** Patterns of rearranged gene orders of Amphibians.

| WANCY | n | IQM | n | HSL | n |
| --- | --- | --- | --- | --- | --- |
| 'W', 'N', 'A', 'C', 'Y'* | 4 | NA | 0 | NA | 0 |
| 'W', 'A', 'C', 'Y', 'N'* | 2 |  |  |  |  |
| 'A', 'C', 'W', 'N', 'Y'* | 3 |  |  |  |  |
| 'W', 'N', 'C', 'Y', 'A' | 1 |  |  |  |  |
| 'W', 'A', 'Y', 'N', 'C' | 1 |  |  |  |  |
| 'N', 'C', 'Y', 'W', 'A' | 1 |  |  |  |  |
| **Total** | 12 |  | 0 |  | 0 |

(\*convergent gene order)

**Table 4.**
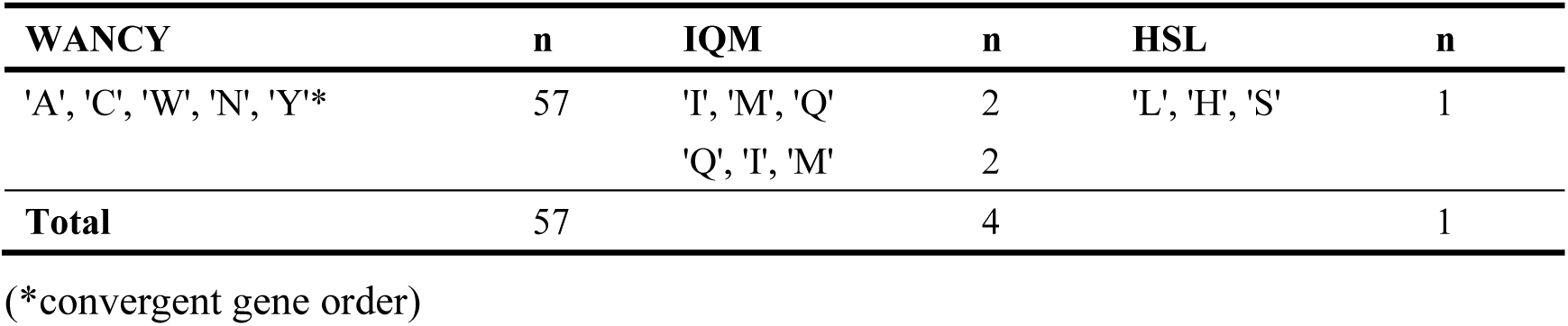
Patterns of rearranged gene orders of Mammalia.

**Table 5.**
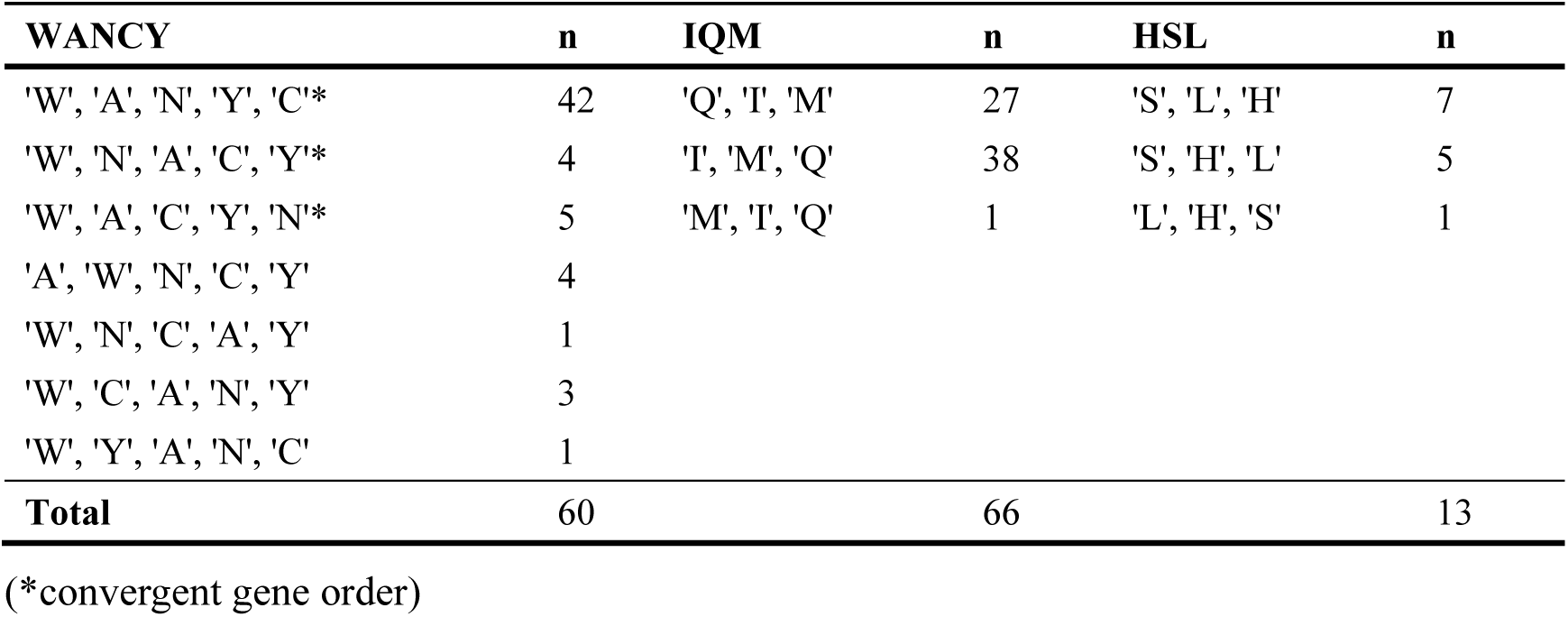
Patterns of rearranged gene orders of Osteichthyes.

**Table 6.**
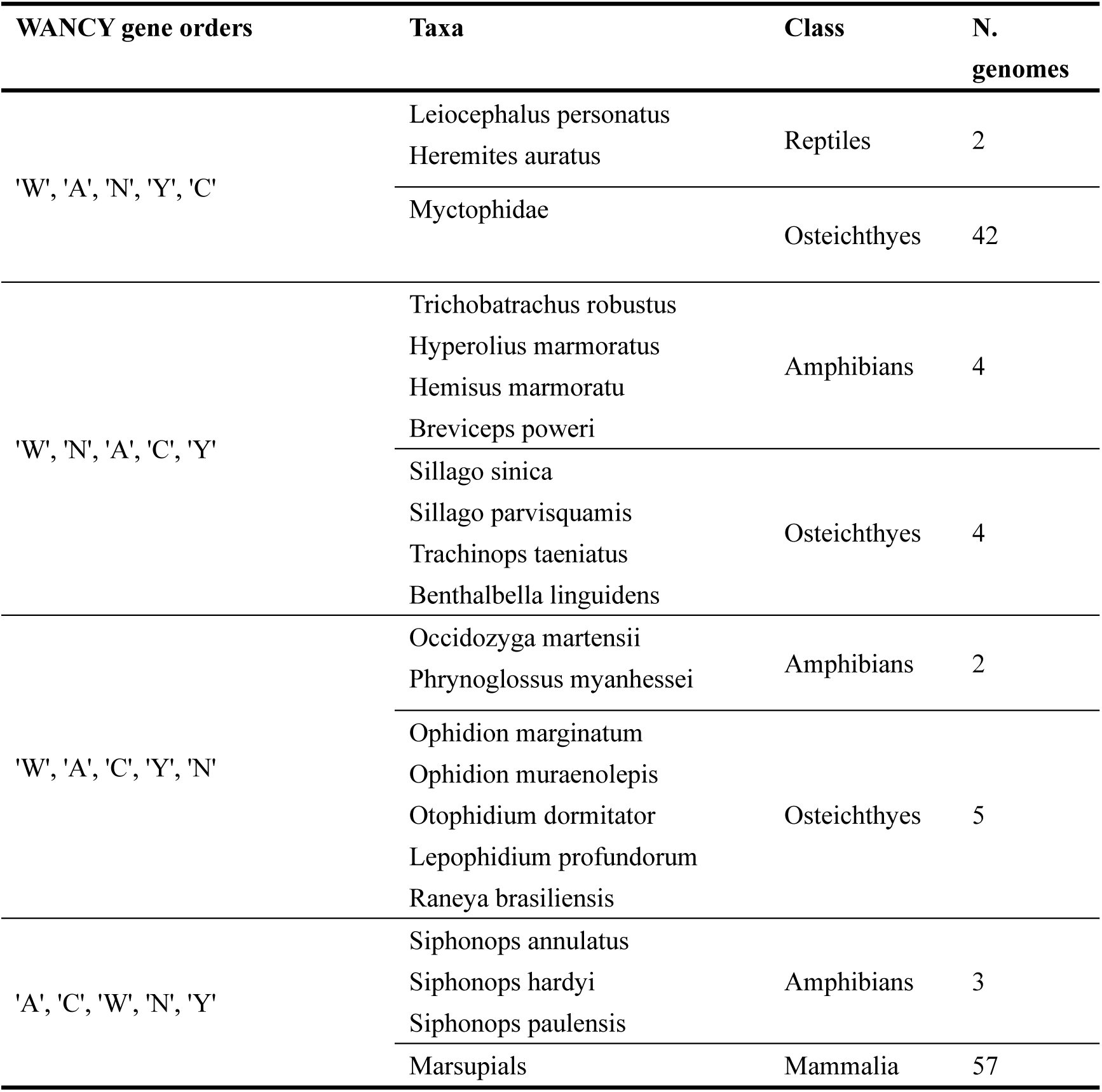
Convergent gene orders in the WANCY region.

Among Reptiles, Amphibians, Mammalia and Osteichthyes, the number of gene order rearrangements in the WANCY region are found to be 2, 12, 57 and 60 respectively. For the IQM region, the numbers are 55, 0, 4 and 66 respectively. For the HSL region, the numbers are 23, 0, 1 and 13 respectively. However, these numbers are not reflective to independent gene order rearrangement events. For a genomic section of five consecutive genes (the WANCY region), the possibility of two different species independently evolved the same genotype can be calculated using the following fomula:

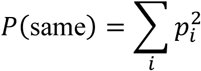

For the five genes, there are 2^5^ = 32 possible copy-retention patterns, of which 6 still yield the original order WANCY, while the remaining 26 each produce a distinct rearranged order. Therefore:

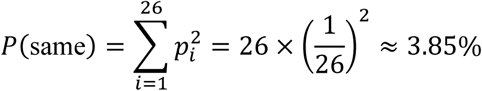

The probability (3.85%) is relatively low, and will continue to drop with the increase in the number of species. For a genomic section of three consecutive genes, the possibility increases to 25%. From another perspective, the frequency of a rearranged gene order which different from the vertebrate consensus for a region of three genes should be 1-(3+1)/2^3^= 50%. However, it can be seen from table 2, among the 55 rearrangements in the IQM region, 54 showed the same gene order of ‘Q’, ‘I’, ‘M’. The frequency of showing the gene order of QIM has reached 98%. This indicates that this gene order may be an ancestral gene order and has been retained in multiple descendant lineages. Therefore, phylogenetic filtering was conducted, the results are shown in the following section.

### 3.2 Calculation of independent rearrangements in the WANCY, IQM and HSL region

Figure 1, 2, 3 and 4 show the phylogenetic filtration of the WANCY rearrangements of Reptiles, Amphibians, Mammalia and Osteichthyes. According to figure 2, for Amphibians, the 11 matched species (not all the rearranged species are yet classified and shown on the tree) show eight independent rearrangement events. And only one of them were TDRL. However, the other seven rearrangement events can also be explained by a single TDRL or a single transposition. That is, in maximum, eight independent TDRL events occurred along the evolutionary path of the matched Amphibians; in minimum, only one TDRL event occurred. However, since not all of the rearranged species were found on the phylogenetic tree (i.e., some species were not yet classified), whether the gene order of these non-match species came from independent rearrangement events needs to be considered separately. For Amphibians, among the three non-match rearranged species, two (*Siphonops annulatus* and *Siphonops hardyi*) belong to the same genus with *Siphonops paulensis*, and they show the same gene order with *Siphonops paulensis* (ACWNY). Therefore, these two rearrangements are not considered as independent events. The other one (*Occidozyga martensii*) shows the same gene order with *Phrynoglossus myanhessei* (WACYN). However, they belong to different families. The rearrangement of *Occidozyga martensii* is therefore recognized as an independent rearrangement event which can be explained by both a single transposition and a single TDRL. Overall, in maximum there were nine independent TDRL events occurred in Amphibians, and one in minimum. Same analyses were applied to Reptiles, Mammalia and Osteichthyes. Results can be seen in the following tables.

**Figure 1.**
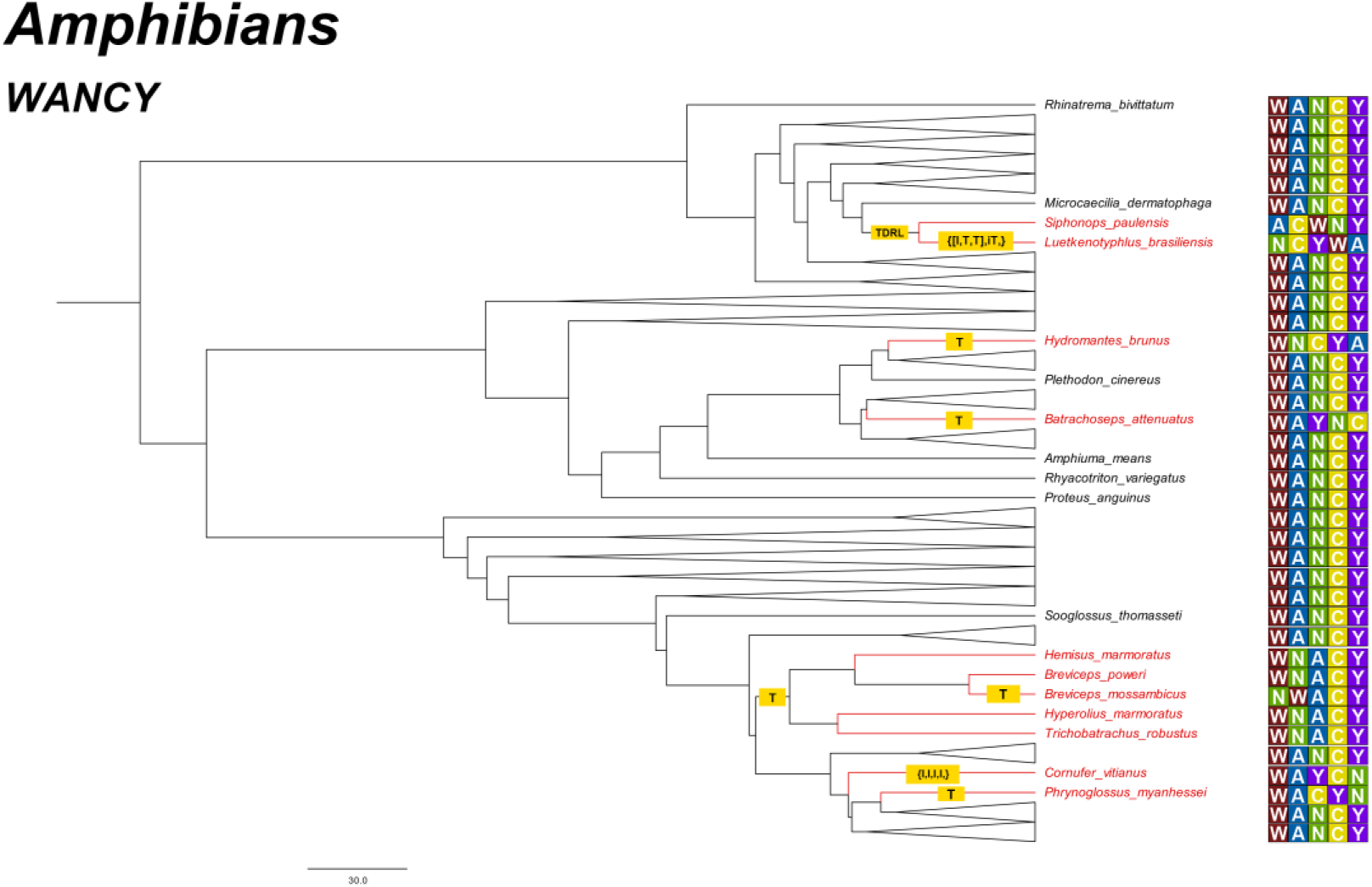
The subtree of Amphibians, demonstrating the evolutionary pathway of rearranged species. The rearrangements on the branches are given as transposition (T), inversion (I), inverse transposition (iT) and tandem-duplication-random-loss events (TDRL). The rearranged species are marked in red. The WANCY gene order of each species was annotated at the right side.

**Figure 2.**
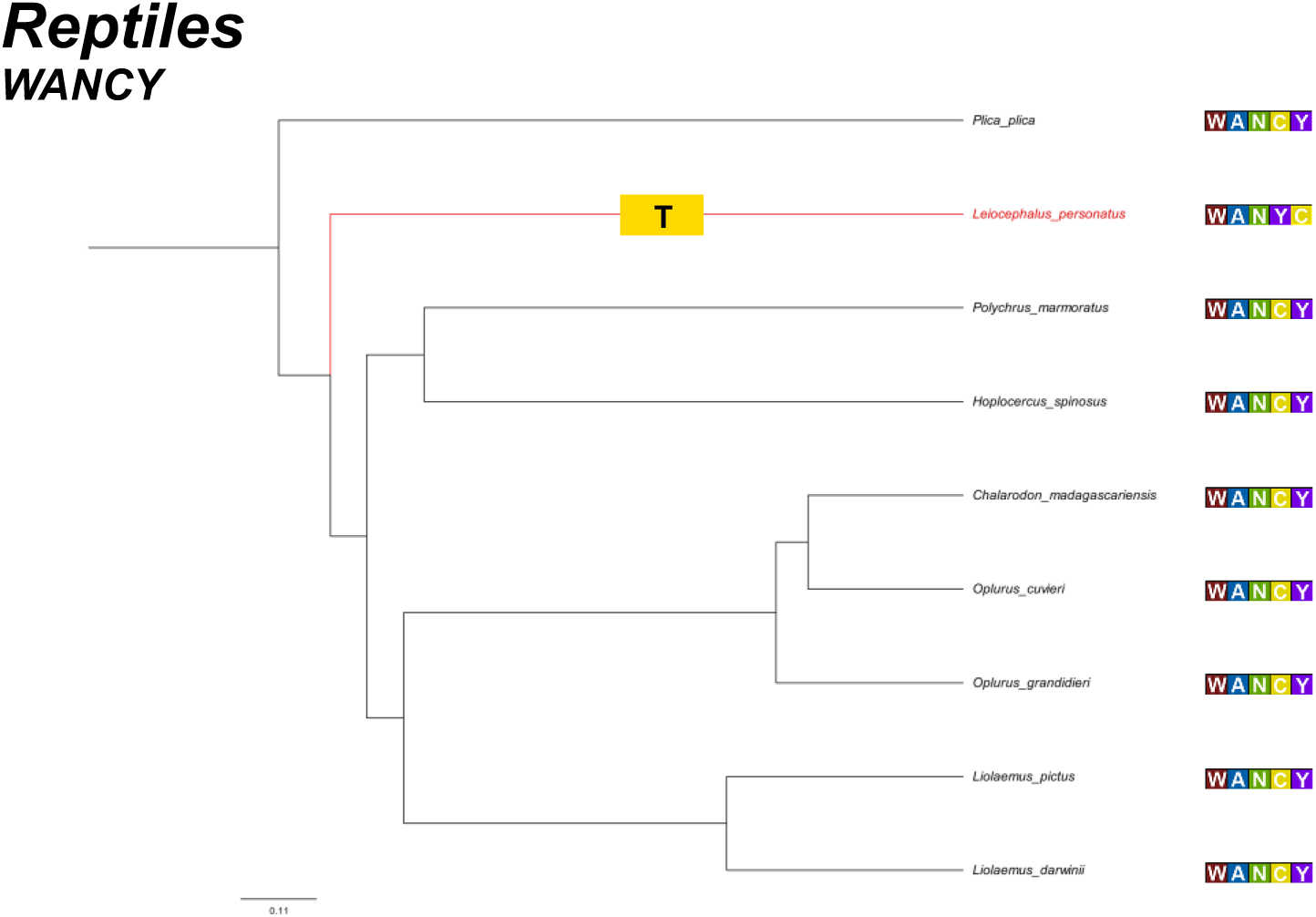
The subtree of Reptiles, demonstrating the evolutionary pathway of rearranged species. Legends see the caption above.

**Figure 3.**
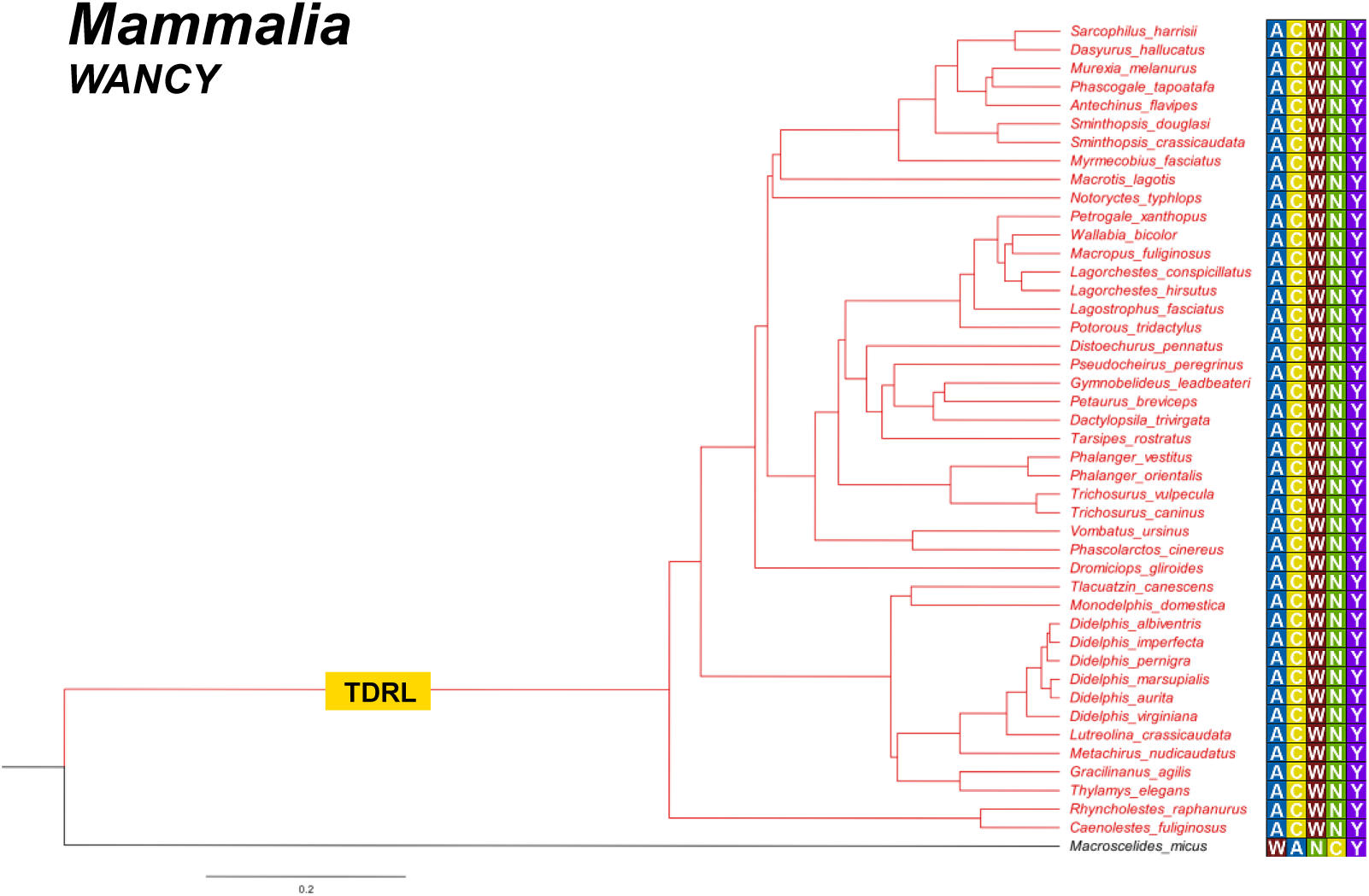
The subtree of Mammalia, demonstrating the evolutionary pathway of rearranged species. Legends see the caption above.

**Figure 4.**
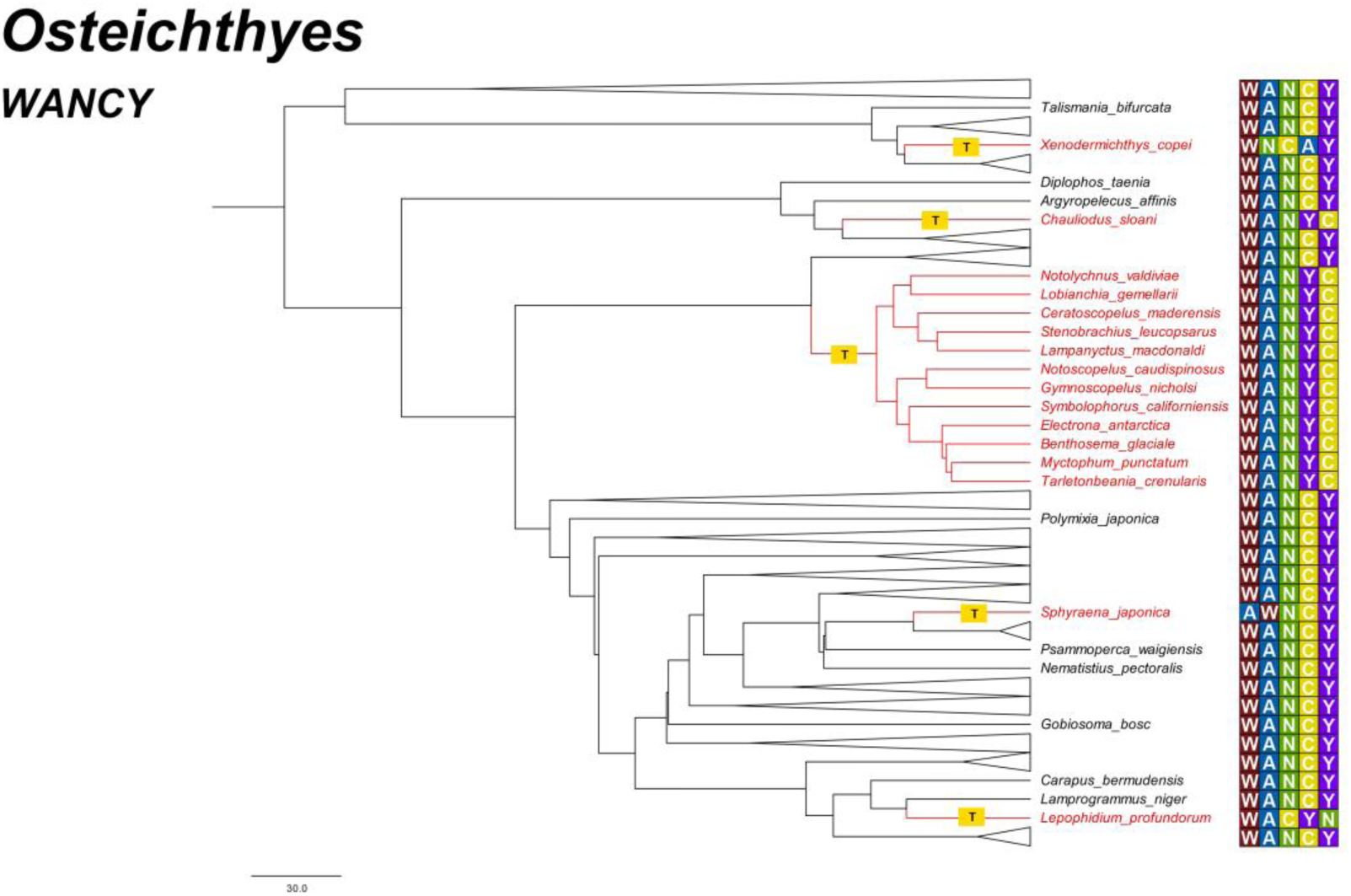
The subtree of Osteichthyes, demonstrating the evolutionary pathway of rearranged species. Legends see the caption above.

According to table 7, when considering the maximum number of TDRL events, Amphibians has the highest probability of occurring TDRL rearrangements (1.85%) among the vertebrates. The probability of occurring TDRL rearrangements in Mammalia, Osteichthyes and Reptiles are 0.06%, 0.26% and 0.30% respectively (0.32% from perspective of vertebrates). According to the percentage of TDRL rearrangements observed in the IQM region, it can be inferred that the probability of TDRL occurrence in the IQM region for Amphibians, Mammalia, Osteichthyes and Reptiles are 0.00%, 0.46%, 0.85% and 1.51% respectively. For the HSL region, the numbers are 0.00%, 0.11%, 0.20% and 0.30% respectively. Eventually, the percentage of TDRL rearrangements in any five consecutive small genes inferred from data of the IQM region for Amphibians, Mammalia, Osteichthyes and Reptiles are 0.00%, 0.37%, 0.69% and 1.23% respectively (0.62% from perspective of vertebrates). The numbers inferred from the HSL data are 0.00%, 0.09%, 0.16% and 0.25% (0.15% from perspective of vertebrates). Binomial test results (table 9) demonstrates that when comparing the WANCY region to the HSL region, the TDRL rearrangement events in the WANCY region is significantly higher (p<0.05). However, when comparing the WANCY region to the IQM region, the TDRL rearrangement events in the IQM region is significantly higher than that in the WANCY region (p<0.01), which shows a completely opposite result. For the minimum TDRL scenario (table 8), two TDRL rearrangement events in the WANCY region were observed among vertebrates, and zero for both the IQM and the HSL region. Although more rearrangement events were observed in the WANCY region, there no significant between the WANCY region and the other two regions on statistics level (p>0.05).

**Table 7.** Quantification of TDRL rearrangements in the WANCY, IQM and HSL regions (All rearrangements which can be explained by both transposition and TDRL are considered TDRL).

|  | Amphibians | Mammalia | Osteichthyes | Reptiles | Total |
| --- | --- | --- | --- | --- | --- |
| **WANCY_TDRL** | 9 | 1 | 13 | 2 | 25 |
| **IQM_TDRL** | 0 | 4 | 21 | 5 | 30 |
| **HSL_TDRL** | 0 | 1 | 5 | 1 | 7 |
| **TOTAL** | 486 | 1751 | 4936 | 661 | 7834 |
| **WANCY_TDRL%** | 1.85% | 0.06% | 0.26% | 0.30% | 0.32% |
| **IQM_TDRL%** | 0.00% | 0.23% | 0.43% | 0.76% | 0.38% |
| **HSL_TDRL%** | 0.00% | 0.06% | 0.10% | 0.15% | 0.09% |
| **TDRL%-IQM** | 0.00% | 0.46% | 0.85% | 1.51% | 0.77% |
| **TDRL%-HSL** | 0.00% | 0.11% | 0.20% | 0.30% | 0.18% |
| **5 genes%-IQM** | 0.00% | 0.37% | 0.69% | 1.23% | 0.62% |
| **5 genes-HSL** | 0.00% | 0.09% | 0.16% | 0.25% | 0.15% |
| **5 genes-re** | 13/16 | 13/16 | 13/16 | 13/16 | 13/16 |
| **3 genes-re** | 0.5 | 0.5 | 0.5 | 0.5 | 0.5 |

**Table 8.** Quantification of TDRL rearrangements in the WANCY, IQM and HSL regions (All rearrangements which can be explained by both transposition and TDRL are considered transposition).

|  | **Amphibians** | **Mammalia** | **Osteichthyes** | **Reptiles** | **Total** |
| --- | --- | --- | --- | --- | --- |
| **WANCY_TDRL** | 1 | 1 | 0 | 0 | 2 |
| **IQM_TDRL** | 0 | 0 | 0 | 0 | 0 |
| **HSL_TDRL** | 0 | 0 | 0 | 0 | 0 |
| **TOTAL** | 486 | 1751 | 4936 | 661 | 7834 |
| **WANCY_TDRL%** | 0.21% | 0.06% | 0.00% | 0.00% | 0.03% |
| **IQM_TDRL%** | 0.00% | 0.00% | 0.00% | 0.00% | 0.00% |
| **HSL_TDRL%** | 0.00% | 0.00% | 0.00% | 0.00% | 0.00% |
| **TDRL%-IQM** | 0.00% | 0.00% | 0.00% | 0.00% | 0.00% |
| **TDRL%-HSL** | 0.00% | 0.00% | 0.00% | 0.00% | 0.00% |
| **5 genes%-IQM** | 0.00% | 0.00% | 0.00% | 0.00% | 0.00% |
| **5 genes-HSL** | 0.00% | 0.00% | 0.00% | 0.00% | 0.00% |
| **5 genes-re** | 13/16 | 13/16 | 13/16 | 13/16 | 13/16 |
| **3 genes-re** | 0.5 | 0.5 | 0.5 | 0.5 | 0.5 |

**Table 9.**
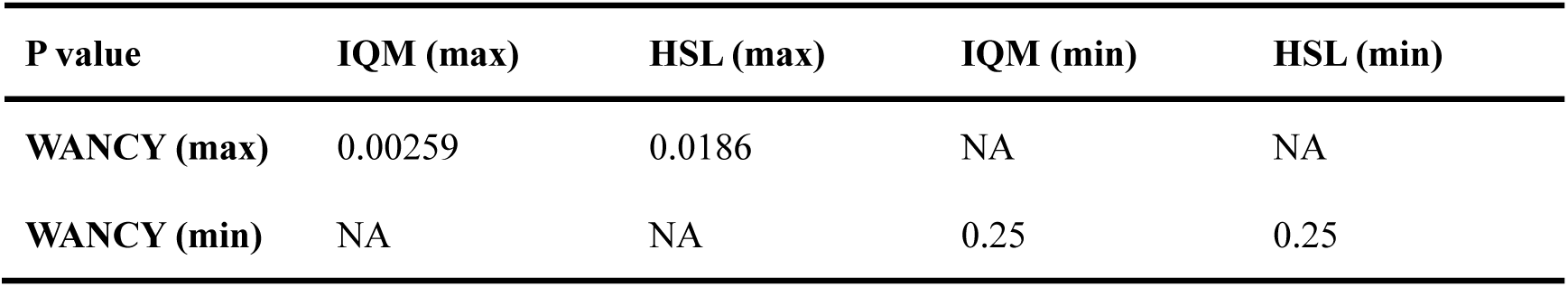
Binomial test results.

## 4. Discussion

### 4.1 Convergent evolution in the WANCY region

The 9419 vertebrate species display 11 rearranged gene orders in the WANCY region. This number is significantly higher than the number shown in the previous study done by San Mauro et al. (2006), in which 453 vertebrate mito genomes displayed four WANCY rearrangements. In this 2006 study, Siphonops species and Marsupials are the only species found to have the same gene order. However, we have now discovered three more convergent gene orders shown in table 6. These convergences may involve different elements of molecular selection processes within the cells (Dubie et al., 2020), or adaptation processes to convergent habitats (Greenway et al., 2020), but the exact reasons are not fully understood.

In general, it is thought that the chances of different taxa sharing the same arrangement of gene order are very low. Mathematically, the 37 standard animal mitochondrial genes can be arranged in a surprisingly large number of permutations if each gene has the same probability of movement. The huge potential number of different permutations implies that the probability of the same gene order occurring in different species lineages is extremely low (Babbucci et al., 2014). However, this study provides further evidence that convergent evolution is a common phenomenon in vertebrate mtDNA rearrangement hotspots. It also supports some previous predictions, e.g., the study of Montaña-Lozano et al. (2022) showed that this convergence is more likely to occur in genes flanking one of the replication origins or in neighbouring tRNA gene segments. This was confirmed by the three gene segments in this study. Another example is Stern (2013) claimed that convergent evolution through homologous site mutations is more common in smaller genomes because there are fewer potential genetic targets that can lead to adaptive phenotypic changes. Our results suggest that the WANCY region convergence coexists in genetically distant orders and relatively close orders and families. This may indicate that there is more convergence of the WANCY regions in small genomes such as the mitochondrial genome than previously thought. The convergence between closer species may be due to the fact that they face similar natural selection pressures. This has been reported in the study of insects, *Locusta migratoria* and *Apis mellifera*, which belong to Orthoptera and Hymenoptera respectively, show the same trnD, trnK versus trnK, trnD arrangement (Flook et al., 1995). In addition to this, this result supports the idea that genes on mito genomes do not have the same likelihood of movement, with tRNAs being the most mobile (Moritz et al. 1987). Different tRNAs move at different speeds, and some of them follow the same paths in different phylogenies, resulting in homogeneous rearrangements (Babbucci et al., 2014). However, the present study only focuses on the hotspot region of the rearrangement. Therefore, it is not capable of indicating whether there is a difference in the convergence frequency between hotspot and non-hotspot regions. In the future, with the support of larger amount of data, the study of convergence on complete small genomes may be able to better reveal the relationship between convergent evolution and mutation targets.

### 4.2 Reasons causing the WANCY hotspot

The results obtained in this study do not fully support the hypothesis that the WANCY region is a mechanistic hotspot. Binomial test presented opposite results when the WANCY was compared with the two regions IQM and HSL. In the scenario where the most TDRLs were considered, the probability of a TDRL rearrangement occurring in the WANCY region was significantly higher than that in any five consecutive tRNA genes calculated from the data of the HSL region. This to a certain extent supports that there is a mechanism in the WANCY region that leads to more TDRLs. At the class level, except for Mammalia in the most TDRL scenario, all the other WANCY regions in both scenarios had more TDRL rearrangements than the rearrangements of five small genes calculated from the HSL data. Therefore, there is a high probability that a mechanistic reason for the WANCY regions exists, regardless of whether the HSL region has a mechanism to promote more rearrangements.

However, the opposite result from the IQM region (significantly more TDRL rearrangements in the IQM than in the WANCY region) does not necessarily refute the hypothesis that the WANCY region is a mechanistic hotspot, since there is no evidence that the IQM is not a mechanistic hotspot. At the same time, we speculate that, comparing to the WANCY region, the gene order rearrangements in the IQM region may be more due to transposition of individual genes than to TDRL. In fact, if we do not consider molecular evidence, all rearrangements of three consecutive small genes can be explained by a single transposition. This has been mentioned in studies on insects, for example all sequenced Ditrysia mtDNAs (Salvato et al., 2008; Timmermans et al., 2014) exhibit trnM translocations on the α-strand, resulting in a different permutation in the IQM region (MIQ). Kong et al. (2009) also reported the transposition of trnQ from light to heavy strand (Q inversion) found in *Cynoglossus semilaevis*. If this is the case, the number of actual TDRL rearrangement events in the IQM region would be much smaller than the current calculations. If more molecular evidence of transposition in the IQM region could be found, it would be more likely to demonstrate that the IQM region has fewer TDRL events than the WANCY region.

Another possible evidence for a mechanistic reason is that we noticed that the rearrangement patterns show that most of the rearranged gene orders in the WANCY region still start with trnW and end with trnY, which means, the actual number of genes involved in the rearrangement maybe less than five. Among the 11 rearranged gene orders, eight of them retained the original position of trnW (72.7%); five of them retained the original position of trnY (45.5%); three of them retained the position of both trnW and trnY (27.3%). If the rearrangements involve fewer genes, then the frequency of actual TDRL events in the WANCY region would continue to increase compared to the present calculations. Consequently, there is a greater tendency to obtain significant results when comparing to the IQM and HSL regions. At the same time, the phenomenon that trnW and trnY remained the same position in most WANCY rearrangements may also indicate that genes trnA, trnN and trnC which closer to the O_L_ are more likely to undergo TDRL and then rearrange. In other words, O_L_ may be the mechanism that promotes the rearrangement of its flanking genes, which in turn leads to the WANCY region becoming a hotspot for TDRL rearrangement. However, another possible explanation for this is that since the trnW gene is located on the heavy strand while the other four genes are on the light strand, considering that it is now generally accepted that slipped-strand mispairing is the main cause of TDRL (Moritz & Brown, 1986), it is reasonable that rearrangements in the WANCY region often do not include trnW. Therefore, more molecular evidence is needed as to whether there is a mechanistic cause for more TDRL in the WANCY region.

The study by Montaña-Lozano et al. (2022) showed that the rearrangements observed in vertebrate mitochondrial genomes are mainly concentrated in certain clusters of tRNA genes, such as trnL1, nad1, trnI, -trnQ, trnM; trnW, -trnA, -trnN, -trnC and trnH, trnS, trnL2, nad5, - nad6, -trnE, cob, trnT, -trnP. These hotspots happen to contain the three regions involved in this study (IQM, WANCY, and HSL). And these three regions are the only gene clusters consisting of three or more tRNA genes on the vertebrate consensus genome. This may imply that the epistemic reason of hotspot formation due to clustering of small genes does exist. However, analyses of reconstructions of historical rearrangement events show that these rearrangements are traced back to only a small number of independent rearrangement events.

For example, the 23 HSL rearrangements shown in the Reptilia are all Crocodilians, and they show the same rearranged order. These rearrangements can be traced back to only one independent ancestral rearrangement event. This is consistent with previously reported results (Janke & Arnason, 1997; Kumazawa & Nishida, 1995) that all crocodilians share the trnS-trnH-trnL gene order (derived from a trnS transposition). The 57 rearrangements in the WANCY region of Mammalia also trace back to only one independent rearrangement event (the Marsupial common ancestor), and the 60 rearrangements in the WANCY region of bony fish can be traced back to up to only 13 independent rearrangement events. I.e., in fact, the HSL region may not be a hotspot for TDRL if independent rearrangement events are used as a criterion for the analysis. The difference in frequency between the WANCY and IQM and the other regions would also be reduced. Consequently, whether an epistemic reason exists remains open to discussion. Actually, it is difficult to prove the existence of an epistemic reason. A more direct way is to prove that the mechanistic reason does not exist in the WANCY region, and then the cause of the hotspot can only be the epistemic reason. Another possible way is to prove that the size of tandem duplications has a distinct distribution over the vertebrate mitochondrial genome, and the peak of this distribution is at least the length of two tRNA genes. If future studies can find this distribution, then it is possible to reveal not only the cause of these rearrangement hotspots, but also a deeper mechanism of tandem duplications.

In addition to the restrictions mentioned above, this study has some other limitations. First, a study of 2831 complete vertebrate mitochondrial genomes by Montaña-Lozano et al. (2022) showed that 20% of the rearrangements detected in these genomic records were due to gene annotation errors. In this study, we were not able to validate gene annotations on all data, which may have led to the identification of some erroneous rearrangements. In future studies, the use of validated gene annotations can yield more accurate statistics when sufficient manpower is available. Second, stronger evidence is missing for the analysis of independent rearrangement events, both for already classified species and for unclassified but rearranged species (not appearing on the phylogenetic tree). Indeed, the lack of molecular evidence leads to the fact that accurately inferring the number of independent rearrangement events is difficult, especially for species that are close together on the phylogenetic tree. For example, in the study of San Mauro et al. (2006), it was reported that although Siphonops and *Luetkenotyphlus brasiliensis* produced different rearranged order, the discovery of the trnN pseudogene proved that the two different sets of rearrangements resulted from the same independent duplication event. That is, the common ancestor of Siphonops and *Luetkenotyphlus brasiliensis* produced two different random losses after a tandem duplication event, which is different from what the TreeREx algorithm had inferred. It was also reported in the study of Xia et al. (2016) that the discovery of intermediate states of functional gene copies reveals that multiple parallel random losses in the single species *Quasipaa Boulengeri* sharing the same tandem duplication event. Admittedly, molecular evidence for TDRL is rare (San Mauro et al., 2006; Hartmann et al., 2016). Due to strong selection pressure, most redundant genes result from tandem duplication are rapidly discarded (San Mauro et al., 2006). Therefore, it is currently difficult to find sufficient evidence to make a convincing speculation on independent rearrangement events. Finally, although this study analysed almost all the available data on GenBank, the results show that, the current volume of data is still insufficient to draw statistically compelling conclusions. Despite the explosion of molecular data today, overall, mitochondrial gene rearrangement remains a rare event. In this study, the analysis of more than 40,000 records of vertebrate mitochondrial genomes yielded only a small number of rearrangement events. In the future, more molecular evidence and more gene sequence data from different species may be able to shed more light on the causes of the WANCY hotspot, both in terms of molecular mechanisms and statistics.

## Conclusion

This summative study on a large number of vertebrates mitochondrial gene data records from NCBI GenBank provides new insights into the patterns and mechanisms on gene order rearrangement. Our study reports three new convergent evolutionary gene orders in the WANCY region, which offers further evidence on the universality of convergent evolution in rearrangement hotspots. With the support a larger amount of data in the future, studies on the whole genome may be able to reveal deeper molecular mechanistic of convergent evolution. In addition to this, the comparison of gene order rearrangement frequency between the WANCY region and the other two similar small gene clusters demonstrated opposite results. Therefore, it is hard to confidently prove the hypothesis that the WANCY region is a mechanistic hotspot from statistical perspective. However, the fact that genes more adjacent to O_L_ are more likely to rearrange to a certain extent supports the possibility of O_L_ being the mechanistic reason which promotes additional rearrangements. Comparing to mechanistic reasons, it is more arduous to prove the existence of epistemic reason. In the future, studies on the distribution of tandem duplication size have the potential to reveal whether epistemic reason contributes to hotspots. In addition, collection of molecular evidence and more gene order records of different species are crucial. The former helps to determine the number of historical independent rearrangement events more accurately, the latter contributes to more significant results on the statistics level. These are essential to reveal the formation of hotspots of gene order rearrangement on mitochondrial genomes of vertebrates from a more profound perspective.

