## Supplementary material for "Statistical analysis provides new insights into patterns and mechanisms of the WANCY hotspot on vertebrate mitochondrial genomes": SI_1. Resampling script.txt

import random

def process_file_with_full_species_name(file_path):
    with open(file_path, 'r') as file:
        lines = file.readlines()

    species_dict = {}
    for line in lines:
        parts = line.strip().split()
        # Full species name is the two elements following the 'bp' text
        if 'bp' in parts:
            bp_index = parts.index('bp')
            if bp_index + 2 < len(parts):
                species_name = parts[bp_index + 1] + ' ' + parts[bp_index + 2]
                species_dict.setdefault(species_name, []).append(line.strip())

    # Selecting a random entry for each species
    selected_entries = [random.choice(entries) for entries in species_dict.values()]

    return selected_entries

file_path = 'C:/Users/xucho/Desktop/Intra TDRL attachments/12864_2016_3309_MOESM6_ESM/mtGord V0.5/geneord-Chondrichthyes.txt'

# Process the file with the revised function for full species name
full_species_processed_entries = process_file_with_full_species_name(file_path)


# save to a new txt file
output_file_path = 'C:/Users/xucho/Desktop/Intra TDRL attachments/12864_2016_3309_MOESM6_ESM/mtGord V0.5/Re-sampled/Chondrichthyes.txt'  

with open(output_file_path, 'w') as file:
    for entry in full_species_processed_entries:
        file.write(entry + '\n')  
