## Supplementary material for "Statistical analysis provides new insights into patterns and mechanisms of the WANCY hotspot on vertebrate mitochondrial genomes": SI_2. WANCY_Rearrangement identification.txt

def check_sequence(line, sequence):
    """
    Check if the specified order is in the given row and the order is inconsistent with the given order.
    """
    for i in range(len(line)):
        if line[i:i+len(sequence)] == sequence:
            return False  # Specified order in the row，but no rearrangement
        if all(gene in line[i:i+len(sequence)] for gene in sequence):
            return True  # Specified order in the row, and rearranged
    return False  # Specified order not in the row

def sequence_present(line, sequence):
    """
    Check if the specified order is in the given row
    """
    for i in range(len(line)):
        if line[i:i+len(sequence)] == sequence:
            return True
    return False

def process_file(input_filename, output_filename):
    """
    Read the iuput file, indentify and extract the individuals with a WANCY rearrangement.
    """
    sequence_5 = ["Trp", "Ala", "Asn", "Cys", "Tyr"]
    sequence_6 = ["Trp", "Ala", "Asn", "RO", "Cys", "Tyr"]
    exclude_sequence_1 = ["Tyr", "Cys", "RO", "Asn", "Ala", "Trp"]
    exclude_sequence_2 = ["Tyr", "Cys", "Asn", "Ala", "Trp"]
    
    """Define 'exclude_sequence' is because some of the records on GenBank are in the reversed order."""

    with open(input_filename, 'r') as infile, open(output_filename, 'w') as outfile:
        for line in infile:
            genes = line.strip().split()
            if (check_sequence(genes, sequence_5) or check_sequence(genes, sequence_6)) and not (sequence_present(genes, exclude_sequence_1) or sequence_present(genes, exclude_sequence_2)):
                outfile.write(line)


input_filename = 'C:/Users/xucho/Desktop/Intra TDRL attachments/12864_2016_3309_MOESM6_ESM/mtGord V0.5/Re-sampled/birds.txt'
output_filename = 'C:/Users/xucho/Desktop/Intra TDRL attachments/12864_2016_3309_MOESM6_ESM/mtGord V0.5/Re-sampled/Re-WANCY/birds.txt'
process_file(input_filename, output_filename)
