## Supplementary material for "Statistical analysis provides new insights into patterns and mechanisms of the WANCY hotspot on vertebrate mitochondrial genomes": SI_5. Interpretation.txt

#For the "Resampling script":

1. The script is based on Pyhton 3.11.
2. The "Resampling script" resamples the gene information records. If there are multiple records of the same species, only one will be randomly retained and the rests will be deleted, to ensure that each species occurs only once.
3. The input file should be a txt file generated by the Perl script "mtGordV0.5.pl" or in the same format. This script was written by Yuchi Zheng of Chengdu Institute of Biology, Chinese Academy of Sciences. An example line of the output files is: AB263940	1587 bp	Agama impalearis	cytb	Thr	Pro	3 genes
Items should be separated from each other by a tab. This perl script can be accessed and downloaded from the reference below.

 Xia, Y., Zheng, Y., Murphy, R.W. et al. Intraspecific rearrangement of mitochondrial genome suggests the prevalence of the tandem duplication-random loss (TDLR) mechanism in Quasipaa boulengeri . BMC Genomics 17, 965 (2016). https://doi.org/10.1186/s12864-016-3309-7

5. Remember to change the directory of "file_path" and "output_file_path" to your own working directory.


#For the "WANCY_Rearrangement identification", "IQM_Rearrangement identification" and "HSL_Rearrangement identification":

1. The scripts are based on Pyhton 3.11.
2. These scripts identify and extract the records with gene order rearrangements in the three genetic regions respectively.
3. The input file does not need to strictly follow the above format, but need to be a txt file consisting of lines containing gene orders in annotation. Items should be separated from each other by a tab.
4. An example line of input: Benthosema pterotum Leu	ND1	Ile	Gln	Met	ND2	Trp	Ala	Asn	Tyr	Cys
5. Remember to change the directory of "input_filename" and "output_filename" to your own working directory.
